# Linking movement and dive data to prey distribution models: new insights in foraging behaviour and potential pitfalls of movement analyses

**DOI:** 10.1101/2022.11.17.516808

**Authors:** Katie R.N. Florko, Courtney R. Shuert, William W.L. Cheung, Steven H. Ferguson, Ian D. Jonsen, David A.S. Rosen, U. Rashid Sumaila, Travis C. Tai, David J. Yurkowski, Marie Auger-Méthé

**Author notes:** **Corresponding author**: Katie R.N. Florko, Aquatic Ecosystem Research Laboratory, University of British Columbia, 2202 Main Mall, Vancouver, BC V6T 1Z4.

## Abstract

**Background:** Animal movement data are regularly used to infer foraging behaviour and relationships to environmental characteristics, often to help identify critical habitat. To characterize foraging, movement models make a set of assumptions rooted in theory, for example, time spent foraging in an area increases with higher prey density.

**Methods:** We assessed the validity of these assumptions by associating horizontal movement and diving of satellite-telemetered ringed seals (*Pusa hispida*) – an opportunistic predator – in Hudson Bay, Canada, to modelled prey data and environmental proxies.

**Results:** Modelled prey biomass data performed better than their environmental proxies (e.g., sea surface temperature) for explaining seal movement, however movement was not related to foraging effort. Counter to theory, seals appeared to forage more in areas with relatively lower prey diversity and biomass, potentially due to reduced foraging efficiency in those areas.

**Conclusions:** Our study highlights the need to validate movement analyses with prey data to effectively estimate the relationship between prey availability and foraging behaviour.

## Background

Due to recent advances in biologging technologies and statistical analyses, animal movement data are increasingly used to provide ecological insights and inform conservation and management strategies [1–3]. As resource availability is a fundamental driver of animal behaviour [4], movement modelling can be used to understand relationships between animal behaviour and the heterogeneous landscapes they exploit. For example, putative foraging behaviours identified from tracking data have been used to define the critical foraging habitat of marine mammals [5].

Theories predicting how predators are expected to maximize foraging success and minimize energetic costs are often the basis for movement model assumptions [5]. A particularly important and common assumption in movement modelling is that animals are expected to use area-restricted search (ARS; less-direct movement, higher turning rates, overall lower speeds of travel) in areas of profitable foraging [6,7], and thus studies link time spent in ARS with prey abundance and foraging activity (e.g., [8–10], Fig. 1). For example, in 2021, over half of the papers that used animal movement modelling to infer foraging behaviour assumed that time spent in ARS increased with prey abundance, 78% assumed that foraging effort increased with prey abundance, 85% assumed that time spent in ARS was an indication of foraging effort, and for studies on diving species, 81% assumed that more frequent diving was associated with higher foraging effort (Fig. 2, Literature Review A1). Additionally, 39% of these papers assumed that areas with more ARS behaviour and/or foraging effort were important areas to focus conservation efforts (Fig. 2, Literature Review A1). Some studies have found these assumed relationships realised. For example, reef manta rays (*Manta alfredi*) used ARS in plankton patches [11], and the number of northern elephant seal (*Mirounga angustirostris*) dives was related to the number of prey consumed [12]. While ARS behaviour can be indicative of foraging effort (e.g., searching, capturing, and handling prey [6,7]), movement analyses rarely provide the true profitability (i.e., capture success) of ARS behaviour. This discrepancy may lead researchers to conclude, perhaps mistakenly, that more time spent in ARS in certain areas indicates “better” foraging conditions.

**Figure 1.**
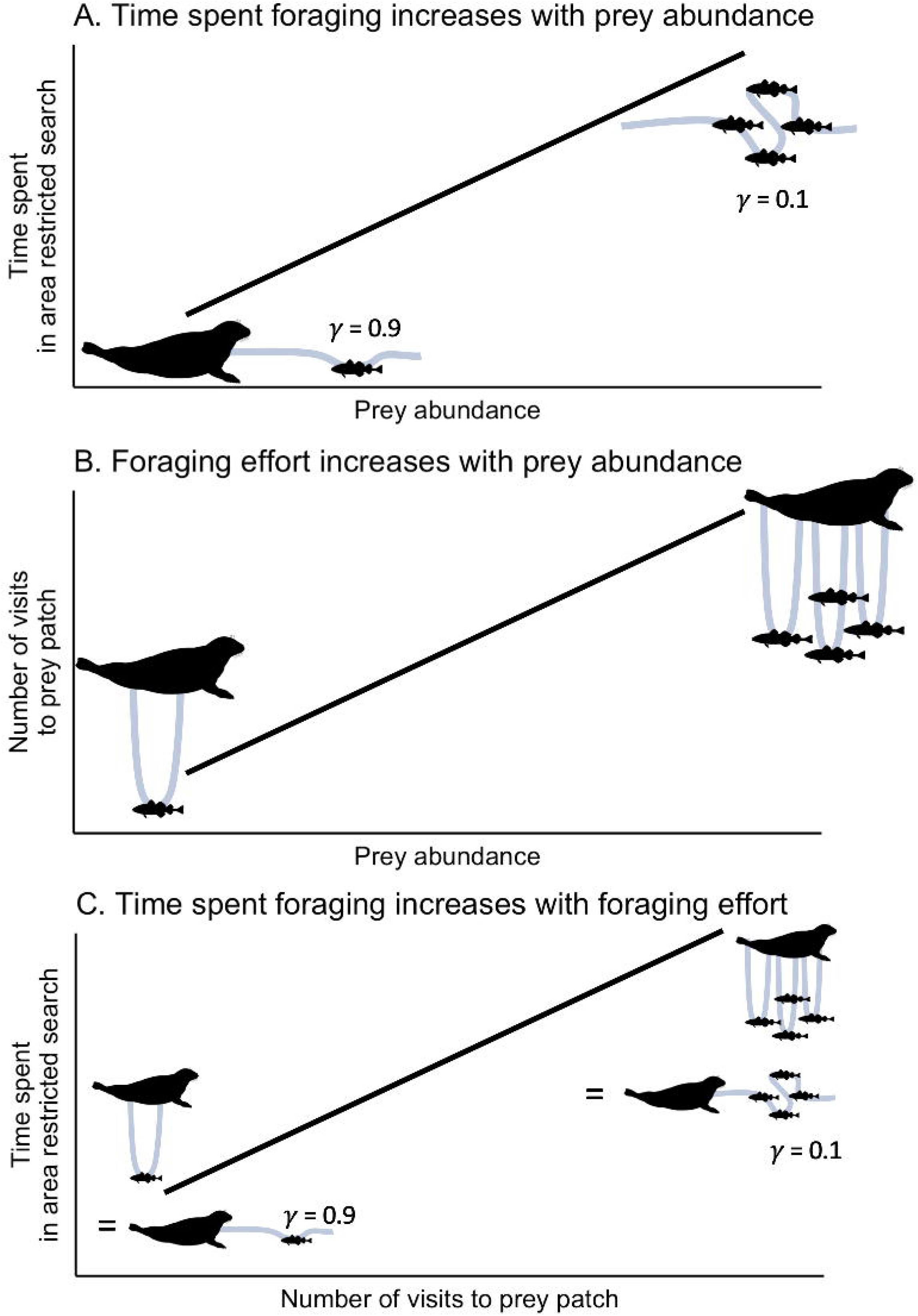
Conceptual plots of three assumed correlations between time spent in an area-restricted search behaviour, foraging effort (e.g., visits to prey patch), and prey abundance. Green lines represent the animal’s trajectory in either A) latitude-longitude movement, B) number of vertical dives, or C) both.

**Figure 2.**
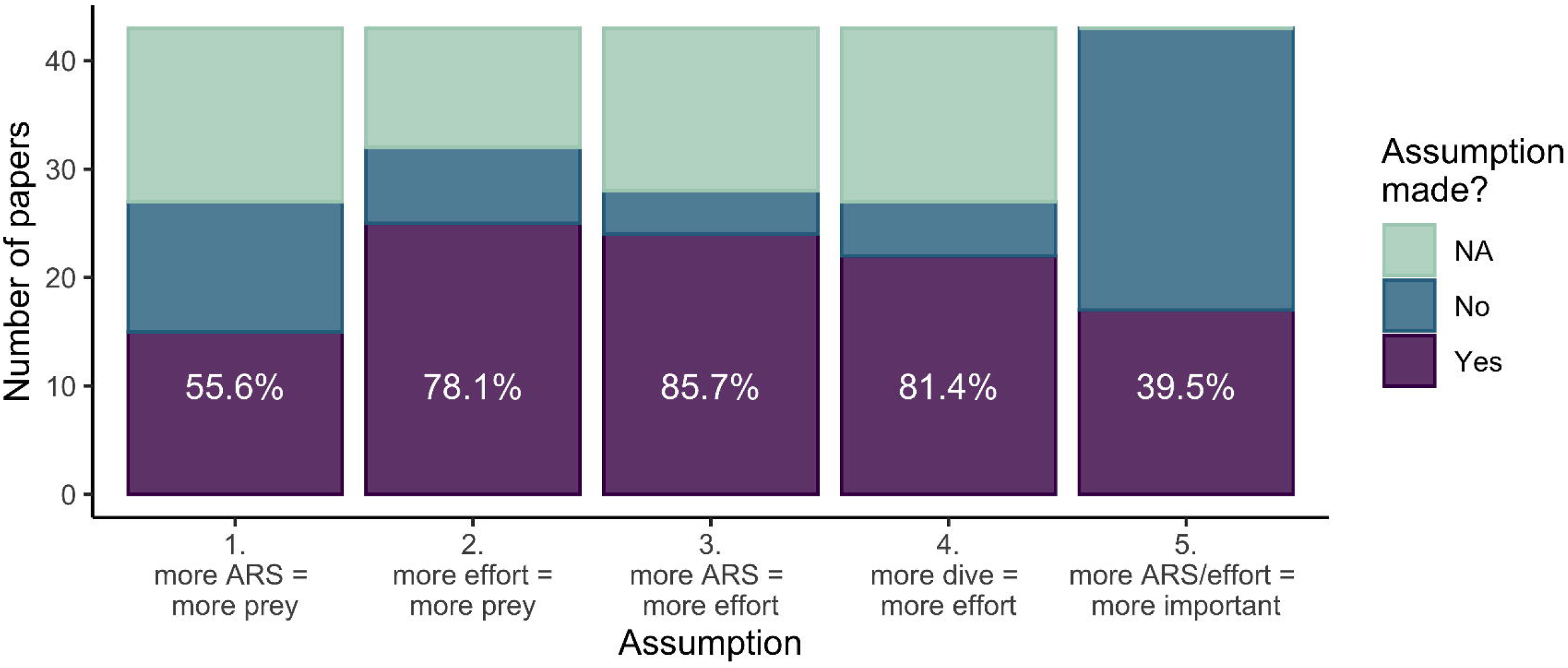
Summary from the systematic review of 43 articles published in 2021 and if they made the following assumptions: Assumption 1: time spent in ARS increased with prey abundance; Assumption 2: foraging effort increased with prey abundance; Assumption 3: time spent in ARS was an indication of foraging effort; Assumption 4: for diving species, more frequent diving was associated with higher foraging effort; and Assumption 5: areas with more foraging (ARS and/or effort such as dives) were important areas for conservation. White text represents the percentage of papers that made the assumption (NAs removed). See Additional File 1 for detailed methodology and results.

While proposed foraging mechanisms, such as ARS, are useful grounds to develop models [7], the complex behaviours of some species can result in unexpected patterns. For example, for generalists, higher prey diversity – independent of prey abundance – may affect foraging effort by increasing the chances of successful prey capture [13], and thus we may expect higher prey diversity to be associated with more time spent foraging. However, contrary to predictions, more time spent foraging in certain areas may indicate poorer foraging conditions (reduced prey density) such as in the case of sink habitats [14]. In addition, animal movement (horizontal and vertical) is influenced by more than just the resource landscape, but also energy, fear, and competition landscapes [15]. Behavioural inference solely based on location data may misrepresent underlying foraging success (e.g. [16]). These nuances can alter the interpretation of animal movement modelling results.

A variety of statistical tools have been developed to infer foraging behaviour from animal tracking data. Quantitative approaches such as state-space models use animal movement (i.e., horizontal location) data to infer behavioural states such as ARS or travel [17–19]. These methods are now able to estimate behaviour as a continuous value of move persistence, on a continuum from 0 to 1, where values towards 0 are indicative of ARS, and values towards 1 are indicative of travel [20,21]. Additionally, newly developed mixed-effects models use environmental covariates to predict behaviour and identify relationships between behaviour and environment [21]. For marine mammals, behavioural estimates may also be inferred by diving data [22] and subsequently, analysis of behavioural data can inform foraging areas (e.g., [23,24]). The development of associated R packages (e.g., the state-space modelling package: foieGras; [25]) have increased the uptake of such analyses.

Results from animal movement models are not typically validated with additional data (but see for example: [26]), which may lead to inaccurate or incomplete interpretation [27,28]. Additionally, analyses often relate the movement of animals to environmental conditions that are proxies for the presence of prey (e.g., [29]), with no a priori hypotheses in terms of the direction of the relationship between foraging behaviour and environmental conditions. Using prey resource data when modelling animal movement allows further investigation of how animals interact with their environment and how we interpret their movement ecology (e.g., [30]). Here, we use movement and diving data from ringed seals (*Pusa hispida*), and modelled prey data, to assess whether: 1) movement models that incorporate information on prey density outperform models using only environmental proxies, and 2) locations with movement and diving behaviours usually classified as foraging are associated with a higher density or diversity of prey.

We analysed the movement and diving ecology of 53 ringed seals with over 14,000 estimated locations (see methods) in Hudson Bay using the recently developed R package mpmm (move-persistence mixed-effects model package; [21]) that incorporated modelled prey biomass estimates [31]. Ringed seals are the most abundant and well-distributed pinniped in the Arctic, where they play an important role in the marine food web as the main conduit of energy between lower trophic levels and top predators (polar bears *Ursus maritimus*) When foraging, seals frequently return to a consistent depth on successive dives [32]. Ringed seals are generalists that forage on locally available prey, including Arctic cod (*Boreogadus saida*), northern sand lance (*Ammodytes dubius*), capelin (*Mallotus villosus*), and various invertebrates [33–35]. Sand lance (a demersal species) is a particularly important prey species for Hudson Bay ringed seals [35]. In years when the proportion of sand lance in their diet is relatively low, their dietary diversity is greater, and their body condition is lower [35]. Estimated ringed seal foraging behaviour has been linked to proxies of resource availability (e.g., chlorophyll-a; [29]), but not in the context of fish biomass estimates through space and time. We used a move-persistence mixed modelling approach to characterize ringed seal behaviour relative to various spatial covariates [21]. Using these modelling results, we 1) ranked models with various estimated prey biomass and diversity variables, and compared their fit to those based solely on environmental variables (e.g., bathymetry), and explored relationships between 2) estimated move-persistence behaviour and foraging effort (i.e., diving), and 3) foraging effort and the estimated prey biomass, diversity, and environmental variables. Our goal was to investigate the nuances of interpretation of emerging and advanced statistical methods for animal movement and provide insights on foraging ecology of ringed seals.

## Methods

### Movement and Dive Data

Ringed seals (n = 53 individuals) were captured in June-November from 2006 to 2012 on the Belcher Islands, Nunavut (Fig. 3). Seals were captured with monofilament mesh nets that were set perpendicular from shore in shallow (<8 m depth) water (full details in [36]). All seals were equipped with ARGOS satellite telemetry transmitters. Specifically, seals captured in the earlier years, 2006-2009 (n = 11), were equipped with SPLASH data loggers (location and time-depth recorders [TDR]) manufactured by Wildlife Computers Ltd (Redmond, Washington, USA). Seals captured in the later years, 2010-2012 (n = 42), were equipped with 9000x data loggers (location and TDR) from the Sea Mammal Research Unit (SMRU, University of St. Andrews, UK).

**Figure 3.**
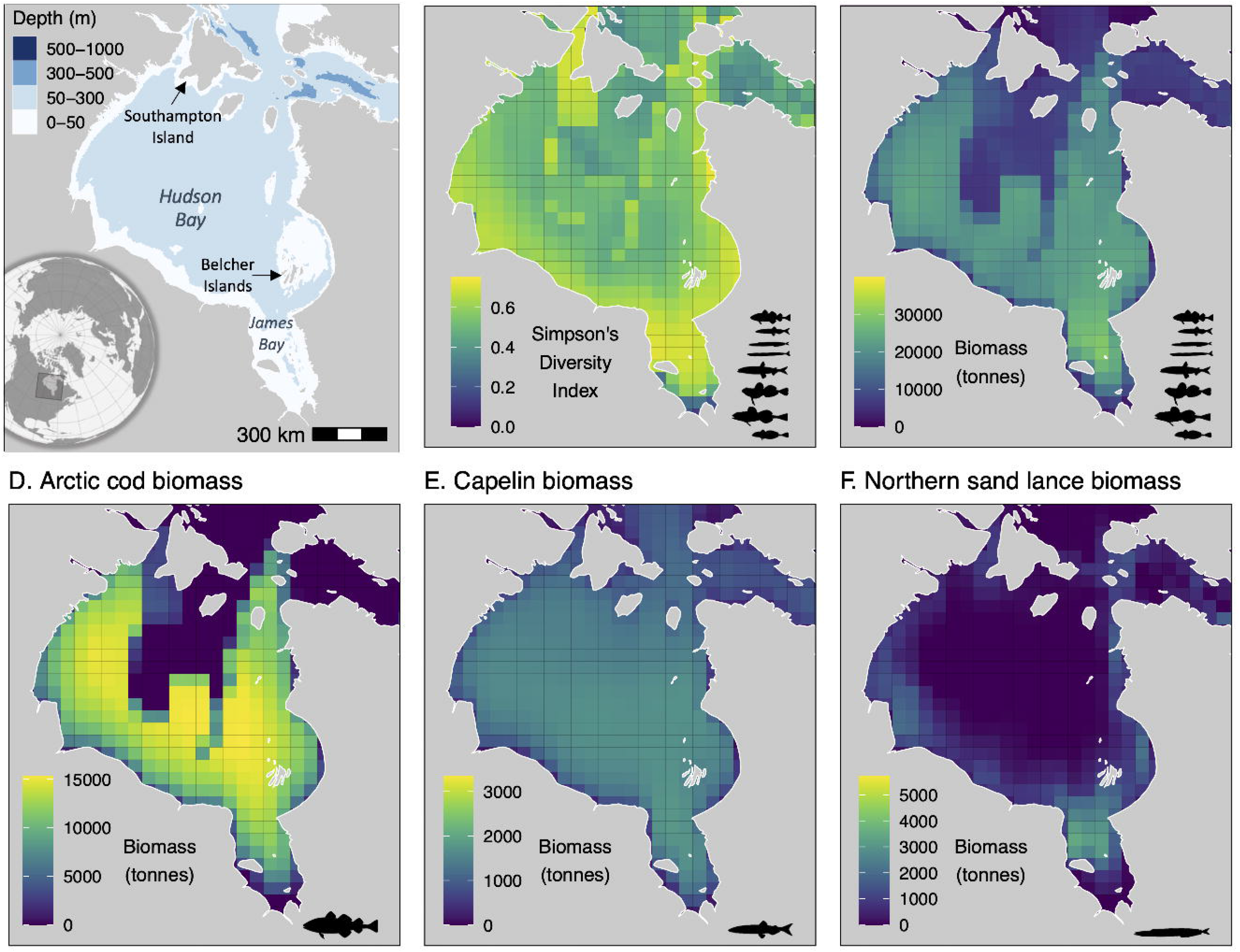
Map of A) study area including bathymetry; B) Simpson’s Diversity Index of 8 fish species; C) total prey biomass of eight fish species; and biomass of D) Arctic cod; E) capelin; and F) northern sand lance. Note difference in scale bars for prey biomass. Prey data summarized from [31]. Data presented in B-F are from 2009, the median year of the study.

We used ringed seal data from the open-water season (i.e., the ice-free summer/autumn) when seals forage intensively to rebuild their depleted energy stores [33], and predation pressure from polar bears is theoretically low. We defined seasons independently for each year [as in 37]. Briefly, we used weekly ice graphs from the Canadian Ice Service (CIS) for the Hudson Bay region. We defined the beginning of the open-water season as the day that sea ice concentration decreased and remained below 50% (i.e., “break-up”) and the end of the open-water season as the day that sea ice concentration increased and remained above 50% (i.e., “freeze-up). We interpolated freeze-up and break-up dates based on the slope of the incline or decline (respectively) between the two weeks where the sea ice threshold was reached.

### Data preparation

We split tracks into multiple smaller segments when transmission halted for >12 hrs and assigned a unique identification number to each segment. We removed tracks with less than 50 transmissions as these led to convergence issues during analysis. This filtering resulted in 124 tracks for the 53 seals (Table A1). The tags transmitted at a frequency of every 1.2 ± 1.7 (mean ± SD) hours for (split) track durations of 19.0 ± 19.3 (mean ± SD) days. Overall, the tags transmitted 41,082 locations.

Since ARGOS location data are observed irregularly in time and are prone to error [38], we filtered and regularized the location data at a 4-hr time step using a correlated random walk state-space model fitted in the foieGras R package [25]. We limited movement rate based on the maximum velocity of ringed seals, ~ 30km/hr. This filtering resulted in 14,639 estimated locations (herein “locations”) to be used in analysis (Table A1).

### Oceanographic data

We extracted the bathymetry (m, 0.01 □ resolution) and monthly sea surface temperature (SST, °C, 0.01 □ resolution) associated with each state-space filtered seal location from the National Oceanic and Atmospheric Administration (NOAA) Environmental Research Division Data Access Program (ERDDAP) data servers from the *etopo180* [39] and *jplMURSST41mday* [40] datasets, respectively, using the rerddapXtracto package [41]. Lower bathymetry values represent deeper depths than higher values.

### Prey data

We used estimated prey biomass data ([31], Fig. 3) from a dynamic bioclimate envelope model, which modelled spatiotemporal changes in the growth, population dynamics, habitat suitability, and movement of each prey fish species from year 1950 to 2100 based on changes in ocean conditions (e.g., sea temperature, pH, salinity) [42,43]. The biomass of each species was modelled at a yearly time step on a 0.5 □ longitude by 0.5 □ latitude grid. The fish data were modelled for both a low-emission (representative concentration pathway, RCP 2.6) and a high-emission (RCP 8.5) scenario [31]; these projections did not diverge during our study period (2006-2013), thus we used the RCP 8.5 data as it aligns more closely to emissions during that time. We matched each seal location in time with the corresponding biomass for each species. We included Arctic cod, capelin, and northern sand lance separately, due to their importance in ringed seal diet [35,44]. We also included the sum of all species (hereby “total prey biomass”), which included Arctic cod, capelin, northern sand lance, as well as Pacific sand lance (*Ammodytes hexaoterus*), Arctic staghorn sculpin (*Gymnocanthus tricuspis*), shorthorn sculpin (*Myoxocephalus Scorpius*), moustache sculpin (*Triglops murrayi*), and rainbow smelt (*Osmerus mordax*). Finally, we calculated Simpson’s diversity index (*D*) among the eight prey species, which reflects the inverse of the probability that two randomly selected prey items are from the same species; lower *D* values are associated with lower species diversity [45]. We calculated *D* as

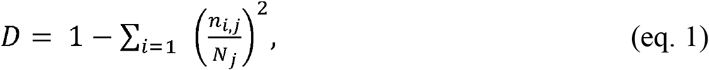

where *n_ij_* = biomass of each species *i* in cell *j*, and *N_j_* = total biomass of all species in each cell *j*, using the R package vegan [46].

### Statistical Analyses

#### Move-persistence mixed-effects models

Move persistence (*γ_t_*; continuous value between 0 and 1) can be used to infer changes in behaviour along animals’ movement paths, where lower values indicate low levels of directional persistence and likely reflects ARS, which is often interpreted as foraging, and higher values indicate high levels of directional persistence, often interpreted as travel [21]. We used a mixed-effects modelling approach to estimate how move persistence varied in relation to bathymetry, SST, and prey biomass and diversity, while incorporating individual variability. We fitted move-persistence mixed-effects models using the mpmm package [21], which models *γ_t_* as a linear function of environmental/habitat predictors,

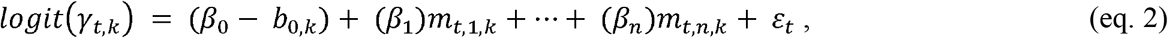

where *β*_0_ is the fixed intercept, *β*_1_,… *β_n_* are the fixed regression coefficients, *m*_*t*,1,*k*_,… *m_t,n,k_* are the predictor variables, *k* indexes individuals, *b*_0,*k*_ is the random deviation for the intercept of individual *k*, and *ε_t_* is the errors where *ε_t_*~*N*(0, *σ_y_*).

#### Model structure and model selection

We fitted the move-persistence mixed-effects models to the state-space filtered and time-regularised seal tracks to infer relationships between behaviour and foraging habitat metrics. Our environmental-only models (proxies for resource availability) included all possible combinations of bathymetry and SST as covariates. Our prey-informed models included models with one of the prey covariates (e.g., capelin biomass), as well as models with bathymetry and each prey covariate together (e.g., bathymetry and capelin) as bathymetry may be more than a proxy for resources and may directly affect the behaviour of seals. All models included random intercepts for individuals, but not random slopes due to convergence issues. All covariates were scaled by year prior to modelling. We fitted the models using maximum likelihood and optimized the models using a bounds-constrained quasi-Newton method (nlminb) and Broyden-Fletcher-Goldfarb-Shanno (BFGS) algorithm. We used Akaike’s information criterion (AIC) to rank models, such that the model with the lowest AIC was characterized as best. If models were within two ΔAIC of the lowest AIC model, we considered the model with the fewest number of estimated parameters as the best model. We used the residuals function in the mpmm package to calculate one-step-ahead residuals from the best model to inspect potential deviations from model assumptions (e.g., normality of the process stochasticity, [47,48]).

#### Leave-one-out cross validation

We assessed the performance of our best-selected model using a leave-one-out cross validation, where we excluded one individual seal to create a new dataset and re-ran the model, and examined the coefficient estimates relative to the full model (all seals), as in [49].

#### Model validation with dive data

To assess whether ARS behaviour was associated with increased dive behaviour, we matched dive data to the move-persistence (γ_t_) values from the best model within two hours of the location data (since movement data was filtered at a four-hour time step). We used the total number of dives and maximum dive depth (i.e., deepest depth during dive), separately, as our dive metrics. We used linear mixed-effects models (LMM) to test the relationship between move persistence and the dive metrics (number of dives, and maximum dive depth, separately) using the nlme package [50]. Additionally, we fitted LMMs to test the relationship between dive metrics and the associated prey data. We fitted the models using maximum likelihood. Each model included track identification number as a random effect, and an AR1 autocorrelation term to account for autocorrelation found in tracking data. We fitted LMMs to test the relationship between dive metrics and associated bathymetry, an important environmental variable, to highlight any differences in consistency relative to the LMMs using prey data. Additionally, we explored additional dive metrics: sum of time spent diving, mean dive depth, cumulative dive depth, mean bottom time, sum of bottom time, and in our bathymetry model: the mean proportion of maximum dive depth of all dives within a two-hour period. Finally, we explored relationships between move persistence from the null model and dive behaviour (see Appendix).

## Results

### Move-persistence mixed-effects models

Ringed seals primarily used the eastern side of Hudson Bay, particularly near the Belcher Islands (Figs. 3, A1), where they were tagged. Seals exhibited a full range of move-persistence values in southeast Hudson Bay (range: 0.01-0.98). Some seals travelled to Southampton Island, western Hudson Bay, and James Bay, and most movements to these locations were more persistent (indicative of traveling), while movement was less persistent (indicative of ARS) near Southampton Island and western Hudson Bay (Figs. 3, A1).

The best-supported model for predicting move persistence included the fixed effects prey diversity and bathymetry (Table 1). Prey diversity was positively associated with move persistence. Bathymetry was negatively related to move persistence, where deeper areas were associated with higher move persistence (Fig. 4, Table A2). Our second-best model included northern sand lance biomass and bathymetry, where northern sand lance biomass was positively associated with move persistence, and again bathymetry was negatively related to move persistence (Table 1, A2).

**Figure 4.**
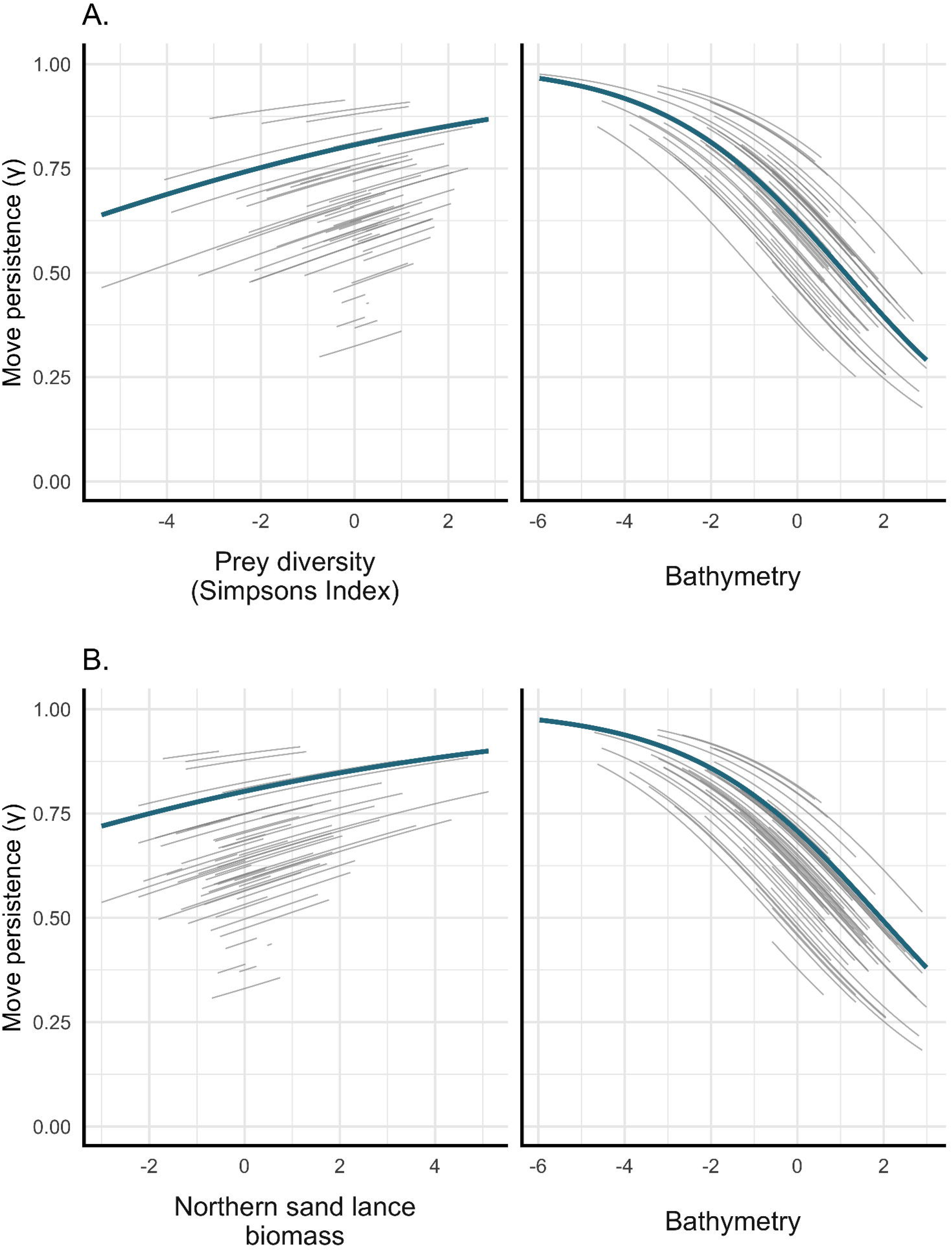
Results from the top two move-persistence mixed model for fixed (thick blue line) and random (individual seals; thin grey lines) effects. The best model, (A), included scaled prey diversity and bathymetry, and the second-best model, (B), included northern sand lance biomass and bathymetry. Low move persistence is indicative of area-restricted search (foraging), and high move persistence is indicative of direct movement (travelling).

**Table 1.** Candidate move-persistence models of ringed seals in Hudson Bay, ranked by Akaike information criterion (AIC).

|  | Model formula | *k* | AIC | ΔAIC |
| --- | --- | --- | --- | --- |
| div2 | ~diversity + bathy + (1 \| id) | 8 | -115846.2 | 0 |
| fish8 | ~biomassnsandlance + bathy + (1 \| id) | 8 | -115844.1 | 2.1 |
| env2 | ~bathy + (1 \| id) | 7 | -115823.6 | 22.6 |
| fish6 | ~biomassarccod + bathy + (1 \| id) | 8 | -115822.5 | 23.7 |
| env1 | ~sst + bathy + (1 \| id) | 8 | -115822.3 | 23.9 |
| fish5 | ~biomassall + bathy + (1 \| id) | 8 | -115822.2 | 24 |
| fish7 | ~biomasscapelin + bathy + (1 \| id) | 8 | -115821.7 | 24.5 |
| fish3 | ~biomasscapelin + (1 \| id) | 7 | -115640.8 | 205.4 |
| fish4 | ~biomassnsandlance + (1 \| id) | 7 | -115640.4 | 205.8 |
| div1 | ~diversity + (1 \| id) | 7 | -115636.4 | 209.8 |
| null1 | ~(1 \| id) | 6 | -115634.8 | 211.4 |
| env3 | ~sst + (1 \| id) | 7 | -115633.9 | 212.3 |
| fish1 | ~biomassall + (1 \| id) | 7 | -115632.8 | 213.4 |
| fish2 | ~biomassarccod + (1 \| id) | 7 | -115632.8 | 213.4 |

*k* = number of parameters estimated; ΔAIC = change in AIC (relative to the best model, div2); diversity = Simpson's Diversity Index of 8 most important prey species; bathy = bathymetry; biomassnsandlance = northern sand lance biomass; sst = sea surface temperature; biomasscapelin = capelin biomass, biomassarccod = Arctic cod biomass; biomassall = biomass of 8 most important prey species.

### Leave-one-out cross validation

The leave-one-out cross validation for our best model indicated that the parameter estimates when one individual was removed were consistently within the confidence intervals of estimates from the model based on all individuals (≥92% within estimate; Table A3). This level of consistency was similar to that of the best model based only on environmental covariates (Table A3). Additionally, the one-step-ahead residuals suggests only minor deviations from model assumptions (Fig. A2, A3).

### Model validation with dive data

We expected a negative relationship between move persistence (from the best model) and all dive metrics. However, we found no significant relationship between move persistence predicted from our best model and number of dives and a positive relationship with maximum dive depth (Table A4, Fig. A4). We also found positive relationships between move persistence and sum of time spent diving, mean dive depth, sum of dive depths, mean bottom time, and sum of bottom time (Table A4, Fig. A4). However, as indicated by the nearly-zero slopes, these relationships are likely a factor of large sample size (Table A4). When using the null model (no covariates) to predict move persistence values, we found similar relationships between move persistence and all dive metrics (Table A4), indicating that these discrepancies are not driven by the specific covariates included in the move persistence model.

We did not find the expected positive relationship between the biomass of bottom dwelling northern sand lance (prey covariate of the second-best model, Table 1) and maximum depth of seal dives (Table A4, Fig. A5). Similarly, we did not find the expected positive relationships between the number of dives (common proxy for foraging effort) and prey (northern sand lance) biomass or prey diversity (Table A4, Figs. A5-A6). However, we found two expected relationships: (1) seals dove deeper in deeper waters than shallow waters (i.e., negative relationship between maximum dive depth and bathymetry); and (2) seals used a greater proportion of the water column in shallow waters (i.e., positive relationship between the proportion of the water column used and bathymetry, Table A4, Fig. A7).

## Discussion

We found that move-persistence models which included prey information outperformed models based solely on environmental proxies of prey distribution, and that prey diversity and bathymetry were the variables that best explained seal movement patterns. We also found that northern sand lance, the most common prey item in the diet of Hudson Bay ringed seals [35], may be an important predictor of seal movement. However, counter to the generally assumed relationship between foraging behaviour and prey density, our results suggest\ that foraging effort – measured as low move persistence – may occur more in areas characterized by lower estimated prey diversity and northern sand lance biomass. While these prey data are modelled and therefore subject to assumptions (see below), our movement models that predicted these unexpected relationships outperformed the null move-persistence model and models incorporating environmental variables alone. Additionally, we did not find the expected relationships between foraging-like movement and diving behaviours, nor those between dive behaviour and modelled prey biomass. Our observations highlight how these assumptions may not be realistic for opportunistic species such as ringed seals. Interpretation of behavioural movement models, regardless of prey data, may be misleading if used to infer important areas for foraging.

The two top models, which explained the movement data almost equally well (ΔAIC <2), included important prey covariates. We assumed that seals would spend more time foraging in areas with higher prey densities and diversity (Fig. 1A). Our top model suggested prey diversity and bathymetry were the most important covariates for explaining move persistence. The importance of prey diversity was expected given that ringed seals are broadly considered opportunistic (generalist) foragers [44], and have greater niche sizes in areas such as the Hudson Bay, where prey diversity is generally higher than at higher latitudes [51]. The second-best model included a relationship between move persistence and northern sand lance biomass, which was expected due to their importance found in diet studies (e.g., [35]). However, we found opposing relationship to that which we predicted, where foraging behaviour (as defined by low move persistence) was associated with low prey density and diversity. These surprising results may indicate that the common assumption that increased time spent in these areas reflects increased foraging success is incorrect. In fact, foraging may conclude earlier in high prey diversity and density areas due to animals swiftly completing sufficient prey capture events (e.g., reached stomach capacity). That is, predators may spend less time foraging in high prey density (or diversity) areas due to high foraging success and more time searching for and capturing prey (i.e., ARS behaviour) in areas with lower prey density and/or patchy environments [52,53]. Specifically, it is possible that the ringed seals in our study sometimes found sufficient prey mid-dive, especially given that the prey densities encountered appeared moderate to high, rather than low (zero) to high (Fig. A8). Alternatively, while ringed seals are considered generalists at the population level, they may also be more specialized at the individual level [51]. Increased prey diversity may lead to a reduction in preferred prey species/type for some individuals, and thus a reduction in foraging behaviour. Additionally, prey species such as sand lance may require more searching due to their small body size and burrowing behaviour, which may explain why more time spent foraging was related to low biomass.

Our study demonstrates how using modelled prey fields, rather than simpler prey proxies, can help avoid inaccurate interpretation of ARS behaviour and aligns with other studies that have shown deviations from the common, and sometimes adequate, assumption that animals use ARS when foraging. ARS has been linked to prey capture for species foraging on patchily-distributed prey, for example, as confirmed from the clicking-behaviour of dolphins [10]. In contrast, low move persistence was related to less intense foraging activity of Adélie penguins (*Pygoscelis adeliae*), opposite to expectations and possibly indicative of resting behaviour [49]. Similarly, depth-accelerometers revealed falsely-identified ARS behaviour when masked boobies (*Sula dactylatra*) were resting on the water’s surface [16]. Additionally, some species may move quickly through foraging areas while still completing successful prey captures (e.g., southern elephant seals, *Mirounga leonina*, [54]).

While we detected the logical relationship that deeper dives were associated with deeper depth, we did not find meaningful and expected relationships between most dive characteristics and apparent foraging effort. While seals may “give up” on a dive and return to the surface if prey density is insufficient [55], they are also expected to spend time searching for and pursuing prey if prey fields are moderate. In these cases, we would expect to see increased dive duration in situations where prey density/diversity was lower, which we also found were associated with lower move persistence. However, we did not find a relationship between move persistence and number of dives, and no dive characteristic was significantly predictive of move persistence. Consistent with [29], we found that ringed seals had low move persistence at shallower depths, whereas more time was spent travelling (high move persistence) at deeper depths.

Modelling is, by definition, a simplistic representation of a complex system. Analyzing animal distribution data using modelled prey data is a valuable exercise that is becoming more common with emerging technology, remote sensing, and modelled data (e.g., [56]), but it is subject to the validity of underlying assumptions. Therefore, it is important to consider whether our results may have been unduly affected by the necessary limitations and assumptions. Our modelling approach did not include invertebrates, which have been found in ringed seal diet analyses, particularly during the spring (found in 15% of ringed seal stomachs; [35]). However, the anticipated relationship where more invertebrate prey would yield lower move-persistence behaviour in seals would require an inverse relationship in distribution between fish and invertebrate biomass. While invertebrate data are not available for Hudson Bay, it is unlikely that their distribution is opposite to that of fish, as we expect areas of high production to be associated with high invertebrate and total estimated forage fish biomass (i.e., along the coastlines, Fig. 3). Our prey dataset was not measured empirically, rather modelled at a relatively coarse spatial resolution (0.5 □ latitude x 0.5 □ longitude grid) which provided insight on the relative distributions throughout Hudson Bay. The heterogeneity among these grid cells may not be meaningful at this spatial scale and thus our move-persistence results may be an artifact of the dynamic bioclimate envelope model. However, in support of these prey density models, the finer spatial scale of sea surface temperature (0.2 □ latitude x 0.2 □ longitude grid), a known predictor of seal movement behaviour [29], did not serve as a better predictor of movement than our modelled prey data. We also found that our movement models that included the prey variables outperformed not only the null model, but also the environment-only models (Table 1). At the broad spatial scale, the seals also appeared to use areas with higher sand lance density (Fig. A8). Additionally, move persistence estimated from our null model (i.e., with no predictor variables) was also not related to the diving metrics, further illustrating the mismatch between our assumptions and results regardless of prey data (Table A4). During our leave-one-out exercise, we found that the prey variables performed approximately as well as environmental variables, which may be expected as prey was modelled using environmental variables [31].

Further, we used AIC to rank our move-persistence models. AIC is a measure of relative model quality, not a measure of absolute model fit [57]. However, our move-persistence models that included modelled prey performed better than the typically-used models which only incorporate prey proxies, as well as the null model, which suggests that modelled prey *relatively* improved model fit.

While recent advances in state-space models have moved beyond the discrete behavioural prediction (i.e., 0 = foraging or 1 = travelling), interpretation of the results remains difficult. ARS behaviour is often assumed to be a single behavioural state – foraging – but may actually be a composite of other behaviours which may not be directly correlated to foraging effort or success [58]. For example, seals commonly rest or sleep at the surface (“bob”) in the water, which would likely result in low move persistence, or may travel at depth [59], resulting in high move persistence, further challenging interpretation. Additional validation using multiple data sources (e.g., jaw accelerometers, [60]; video recorders, [26]) may help to determine how accurately movement data can be used to classify foraging behaviour for further model refinement. Additionally, identifying relationships between prey distribution and predator behaviour may be further complicated by spatiotemporal scale, where at a certain (unknown) scale it is difficult to differentiate between foraging and travelling (for example, as studied in wandering albatross (*Diomedea exulans*): [61], elk (*Cervus canadensis*): [62], and reef manta rays: [11]). We regularised our movement data at a four-hour time step, which may represent a short duration relative to other studies (e.g., [21] used a 24-hour time step) but a long duration for foraging and movement ecology. Choosing a relevant species-specific time step that reflects behavioural “states” should not be too fine-scale (i.e., detecting behavioural “events”) or too coarse (i.e., combining multiple behaviours; [27]) and may require additional field observations and technologies (i.e., animal-borne video cameras) to understand the durations of certain behaviours for better interpretation.

Fear of predators, and inter- and intra-specific competition affects the behaviour, foraging patterns, and distribution of prey species [63,64]. Prey may avoid regions where perceived predation risk is high and forgo feeding opportunities in order to reduce predation risk, as has been found with kangaroo rats (*Dipodomys merriami*, [65]). Ringed seals may experience fear throughout Hudson Bay from polar bears and possibly killer whales (*Orcinus orca*, [66]). Seal prey biomass may be relatively high near the shoreline, but this habitat may be associated with increased polar bear abundance and therefore “riskier” habitat. Ringed seals are also harvested by subsistence hunters and may perceive habitats near communities as riskier. We focused our study on ringed seals during the summer/autumn period when predation pressures from polar bears are removed, but seals may still exhibit antipredator behaviours (e.g., vigilance in lieu of foraging), which may contribute to additional noise in the movement and diving data [67]. Additionally, harbour seals (*Phoca vitulina*) and harp seals (*Pagophilus groenlandicus*) occur in Hudson Bay and share a similar diet with ringed seals [68,69] and thus may contribute to interspecific competition. Intraspecific habitat segregation amongst age classes is known to occur in many pinnipeds (e.g., [70]). Adult ringed seals forage under land-fast ice and subadults forage further offshore during the ice-covered winter and spring [33], which may be related to intraspecific competition.

## Conclusions

Our study demonstrated that although new tools to estimate drivers of animal movement may suggest important relationships between habitat and behaviour in some species, these relationships need to be considered carefully for opportunistic species, or species that have a clear disconnect between horizontal movement and diving activity. There are many nuances to interpreting results from animal movement and associated behavioural estimates, but these models are appropriate for testing ecological hypotheses on the areas and covariates associated with where animals spend relatively more time exhibiting various behaviours. While we have explored reasons why more foraging-like behavior may not indicate better foraging habitat (e.g., low modelled prey density and/or foraging success), areas where animals are spending relatively more time are still important areas for habitat protection and conservation [2]. Additionally, incorporation of multiple data types as a validation exercise can provide essential insight on presumed behaviours. Our work highlights that interpreting movement behaviour results may be precarious without prior knowledge on prey availability, and that caution in providing interpretations is warranted when this information is not available. These nuances should be considered as statistical methods for animal movement data continue to become more advanced and accessible, and as identifying habitat to protect depends on effective analysis of movement data.

## List of Abbreviations

ARS: area restricted search
TDR: time-depth recorders
SST: sea surface temperature
SMRU: Sea Mammal Research Unit
CIS: Canadian Ice Service
NOAA: National Oceanic and Atmospheric Administration
ERDDAP: Environmental Research Division Data Access Program
RCP: Representative concentration pathway
*D*: Simpson’s Diversity Index
nlminb: Bounds-constrained quasi-Newton method
BFGS: Broyden-Fletcher-Goldfarb-Shanno
AIC: Akaike’s information criterion
(LMM): Linear mixed-effects models

## Ethics approval and consent to participate

All applicable institutional and/or national guidelines for the care and use of animals were followed. Animal handling was approved by the Freshwater Institute Animal Care Committee (FWI-ACC-2006 to 2012) and Fisheries and Oceans Canada’s Licenses to Fish for Scientific Purposes (S-05/06 - 09/13-1006-NU).

## Consent for publication

Not applicable.

## Availability of data and materials

The fish dataset is available in the Dryad repository, https://doi.org/10.5061/dryad.x69p8czjs. The seal dataset supporting the conclusions of this article is available from the corresponding author (KRNF) upon reasonable request.

## Authors’ contributions

KRNF and MAM designed the study, DJY and SHF collected the seal data, TCT and WWLC estimated the fish data, CRS, IDJ, and MAM provided statistical and coding insight, DR provided advice on interpretation, KRNF did the analysis, prepared figures and tables, and wrote the manuscript with contribution from MAM. All authors provided ideas and editorial feedback.

## Acknowledgements

We gratefully acknowledge the Inuit hunters and the Sanikiluaq Hunters and Trappers Association for assistance in the field, especially Lucassie and Johnassie Ippak and their families. We thank B. Hunt and C. Harley for helpful feedback on earlier drafts of this manuscript.

