## Supplementary material for "Linking movement and dive data to prey distribution models: new insights in foraging behaviour and potential pitfalls of movement analyses"

**Additional File 1**

**Literature Review A1: Methods for literature search on assumptions made about animal movement, foraging behaviour, and foraging effort.**

**Literature key word searches:** We conducted a literature search to gather papers that used animal movement modelling to infer foraging behaviour and prey availability. We aimed to assess how common the following assumptions were in the recent literature: 1) time spent in area-restricted search (ARS) increases with prey abundance, 2) foraging effort increases with prey abundance, 3) time spent in ARS is an indication of foraging effort, and 4) for diving species, more frequent diving is associated with higher foraging effort. Further, we assessed if these studies associated areas with more ARS behaviour or foraging effort as areas that were important for conservation and wildlife management. An English language literature search was completed on 29 September 2022 in the Web of Science (WoS; Clarivate Analytics). We searched for peer-reviewed articles published between 1 January 2021 and 31 December 2021. Search terms were entered as topic terms in the article title, abstract, author-specified key words, and WoS-assigned key words. The search included the following search terms:

*(animal movement OR telemetry OR biologging) AND*

*(foraging OR feeding OR prey OR ARS) AND*

*(state-space model OR hidden Markov model OR move-persistence mixed-model OR diving OR dive)*

The search yielded 69 articles. The goal of this literature search was to assess the frequency of these assumptions in biologging studies, and thus we excluded articles that did not use empirical or simulated movement data (i.e., review or methods articles). In total, 43 unique articles were identified and included as part of this review.

**Classification of studies:** For each article, we recorded if the study assumed that 1) time spent in ARS increased with prey abundance, 2) foraging effort increased with prey abundance, 3) time spent in ARS was an indication of foraging effort, and 4) for diving species, more frequent diving was associated with higher foraging effort. Each assumption was classified as ‘yes’, ‘no’, or ‘not applicable’ if they did not include the corresponding data type (e.g., no dive data). Finally, we assessed if the study assumed that areas where area-restricted search was used, or areas with frequent foraging effort (e.g., dives or area-restricted search) were associated with important areas to conserve (‘yes’ or ‘no’). Some studies did not use the term “area-restricted search” but described the concept (e.g., reduced speed, increased turning frequency); these studies were still deemed to consider the concept of ARS and were assessed for assumptions.

**Results:** The 43 unique and peer-reviewed papers that linked animal movement modelling, foraging behaviour, and/or prey availability included studies on a variety of species such as bowhead whales (*Balaena mysticetus*), chinstrap penguins (*Pygoscelis antarcticus*), and harbour seals (*Phoca vitulina*). Of these papers, 55.6% (15/27) assumed that time spent in ARS increased with prey abundance, 78.1% (25/32) assumed that foraging effort increased with prey abundance, 85.7% (24/28) assumed that time spent in ARS was an indication of foraging effort, and 81.4% (22/27) assumed that more diving was associated with higher foraging effort (Fig. 2. Additionally, 39.5% (17/43) of the papers reviewed assumed that areas with more ARS behaviour and/or foraging effort were of conservation importance (Fig. 2).

**Table A1**. Summary of ringed seal movement data used in analysis. Note that this summary represents the data that was used for the foieGras state-space model that regularized (and therefore reduced) the sampling of continuous locations (4-hour time-step).

| **Tag ID (n of batches)** | **Track batch ID** | **Locations (n)** | **First date** | **Last date** | **Duration of track** |
| --- | --- | --- | --- | --- | --- |
| 39384 (1) | 39384_4 | 56 | 2007-09-26 | 2007-10-05 | 9.5 days |
| 39385 (1) | 39385_6 | 100 | 2007-09-17 | 2007-10-05 | 18.5 days |
| 83987 (2) | 83987_3 | 83 | 2008-09-07 | 2008-09-22 | 14.2 days |
|  | 83987_6 | 110 | 2008-09-28 | 2008-10-17 | 19.5 days |
| 94528 (10) | 94528_3 | 56 | 2009-08-12 | 2009-08-16 | 3.4 days |
|  | 94528_5 | 102 | 2009-08-27 | 2009-09-04 | 7.4 days |
|  | 94528_7 | 118 | 2009-09-05 | 2009-09-16 | 10.2 days |
|  | 94528_13 | 101 | 2009-09-26 | 2009-10-04 | 7.7 days |
|  | 94528_17 | 52 | 2009-10-14 | 2009-10-19 | 4.4 days |
|  | 94528_19 | 139 | 2009-10-22 | 2009-11-02 | 10.5 days |
|  | 94528_20 | 119 | 2009-11-02 | 2009-11-12 | 10.3 days |
|  | 94528_21 | 68 | 2009-11-13 | 2009-11-18 | 4.5 days |
|  | 94528_22 | 72 | 2009-11-18 | 2009-11-23 | 5.2 days |
| 94537 (5) | 94537_2 | 84 | 2009-08-09 | 2009-08-14 | 4.8 days |
|  | 94537_3 | 137 | 2009-08-19 | 2009-08-27 | 8.1 days |
|  | 94537_4 | 795 | 2009-08-28 | 2009-10-24 | 57.3 days |
|  | 94537_5 | 69 | 2009-10-25 | 2009-10-29 | 4.0 days |
|  | 94537_6 | 394 | 2009-10-30 | 2009-11-23 | 24.0 days |
| 83985 (1) | 83985_11 | 75 | 2008-10-06 | 2008-10-19 | 12.7 days |
| 43852 (3) | 43852_1 | 86 | 2010-08-22 | 2010-08-28 | 5.3 days |
|  | 43852_4 | 1199 | 2010-09-01 | 2010-11-18 | 78.3 days |
|  | 43852_5 | 87 | 2010-11-19 | 2010-11-23 | 4.6 days |
| 43853 (3) | 43853_1 | 660 | 2010-08-22 | 2010-09-28 | 37.3 days |
|  | 43853_2 | 679 | 2010-09-29 | 2010-11-09 | 41.5 days |
|  | 43853_3 | 233 | 2010-11-10 | 2010-11-23 | 13.5 days |
| 43851 (4) | 43851_1 | 92 | 2010-08-22 | 2010-08-27 | 5.0 days |
|  | 43851_8 | 232 | 2010-09-13 | 2010-09-30 | 17.0 days |
|  | 43851_9 | 733 | 2010-09-30 | 2010-11-12 | 42.9 days |
|  | 43851_13 | 61 | 2010-11-20 | 2010-11-23 | 3.5 days |
| 43837 (2) | 43837_1 | 294 | 2010-08-24 | 2010-09-07 | 14.8 days |
|  | 43837_2 | 1627 | 2010-09-08 | 2010-11-23 | 76.5 days |
| 43845 (2) | 43845_2 | 395 | 2010-08-23 | 2010-09-18 | 26.9 days |
|  | 43845_3 | 951 | 2010-09-19 | 2010-11-23 | 65.5 days |
| 43858 (1) | 43858_1 | 1411 | 2010-08-21 | 2010-11-23 | 94.2 days |
| 43838 (1) | 43838_2 | 1655 | 2010-08-25 | 2010-11-23 | 90.6 days |
| 43836 (2) | 43836_2 | 200 | 2010-08-23 | 2010-09-06 | 13.6 days |
|  | 43836_3 | 1208 | 2010-09-06 | 2010-11-23 | 78.1 days |
| 43857 (1) | 43857_1 | 1794 | 2010-08-21 | 2010-11-23 | 94.7 days |
| 94527 (4) | 94527_2 | 55 | 2009-08-10 | 2009-08-14 | 3.9 days |
|  | 94527_4 | 429 | 2009-08-18 | 2009-09-15 | 28.5 days |
|  | 94527_5 | 280 | 2009-09-16 | 2009-10-03 | 17.1 days |
|  | 94527_6 | 783 | 2009-10-04 | 2009-11-23 | 50.9 days |
| 94526 (4) | 94526_2 | 331 | 2009-08-10 | 2009-08-29 | 18.5 days |
|  | 94526_3 | 652 | 2009-08-30 | 2009-10-04 | 35.0 days |
|  | 94526_5 | 354 | 2009-10-06 | 2009-10-30 | 23.3 days |
|  | 94526_10 | 328 | 2009-11-04 | 2009-11-23 | 19.3 days |
| 43848 (1) | 43848_1 | 195 | 2010-07-09 | 2010-07-17 | 8.0 days |
| 94530 (9) | 94530_1 | 127 | 2009-08-01 | 2009-08-07 | 6.3 days |
|  | 94530_8 | 175 | 2009-08-18 | 2009-08-27 | 9.2 days |
|  | 94530_9 | 66 | 2009-08-28 | 2009-09-01 | 4.4 days |
|  | 94530_13 | 115 | 2009-09-09 | 2009-09-15 | 6.4 days |
|  | 94530_14 | 74 | 2009-09-16 | 2009-09-20 | 4.6 days |
|  | 94530_16 | 439 | 2009-09-22 | 2009-10-14 | 22.8 days |
|  | 94530_17 | 102 | 2009-10-15 | 2009-10-20 | 5.1 days |
|  | 94530_18 | 65 | 2009-10-21 | 2009-10-24 | 3.2 days |
|  | 94530_19 | 649 | 2009-10-25 | 2009-11-23 | 29.9 days |
| 94535 (8) | 94535_7 | 321 | 2009-08-11 | 2009-08-29 | 17.4 days |
|  | 94535_8 | 125 | 2009-08-29 | 2009-09-07 | 8.4 days |
|  | 94535_9 | 146 | 2009-09-07 | 2009-09-14 | 7.4 days |
|  | 94535_10 | 226 | 2009-09-15 | 2009-09-30 | 15.5 days |
|  | 94535_11 | 264 | 2009-10-01 | 2009-10-16 | 15.5 days |
|  | 94535_12 | 306 | 2009-10-17 | 2009-11-05 | 19.5 days |
|  | 94535_13 | 90 | 2009-11-06 | 2009-11-11 | 5.2 days |
|  | 94535_14 | 191 | 2009-11-12 | 2009-11-23 | 10.5 days |
| 94539 (5) | 94539_1 | 132 | 2009-08-01 | 2009-08-09 | 8.6 days |
|  | 94539_2 | 687 | 2009-08-10 | 2009-09-17 | 38.3 days |
|  | 94539_3 | 178 | 2009-09-18 | 2009-09-27 | 9.1 days |
|  | 94539_4 | 455 | 2009-09-28 | 2009-10-19 | 21.7 days |
|  | 94539_5 | 805 | 2009-10-20 | 2009-11-23 | 34.3 days |
| 107827 (3) | 107827_1 | 150 | 2011-06-25 | 2011-07-03 | 7.9 days |
|  | 107827_2 | 77 | 2011-07-04 | 2011-07-09 | 5.0 days |
|  | 107827_6 | 73 | 2011-07-13 | 2011-07-17 | 3.4 days |
| 106386 (2) | 106386_1 | 89 | 2011-06-23 | 2011-06-28 | 5.1 days |
|  | 106386_5 | 190 | 2011-07-04 | 2011-07-14 | 9.4 days |
| 107830 (8) | 107830_1 | 226 | 2011-06-20 | 2011-07-02 | 11.3 days |
|  | 107830_2 | 649 | 2011-07-02 | 2011-07-31 | 28.9 days |
|  | 107830_3 | 54 | 2011-08-01 | 2011-08-02 | 1.6 days |
|  | 107830_12 | 81 | 2011-08-16 | 2011-08-22 | 6.3 days |
|  | 107830_15 | 161 | 2011-08-28 | 2011-09-09 | 11.2 days |
|  | 107830_16 | 348 | 2011-09-10 | 2011-09-30 | 19.6 days |
|  | 107830_17 | 1241 | 2011-09-30 | 2011-11-23 | 54.4 days |
| 106373 (1) | 106373_1 | 406 | 2011-10-30 | 2011-11-23 | 24.4 days |
| 116484 (1) | 116484_1 | 533 | 2012-10-29 | 2012-11-22 | 24.2 days |
| 116485 (1) | 116485_1 | 372 | 2012-10-29 | 2012-11-22 | 24.2 days |
| 116493 (2) | 116493_1 | 195 | 2012-10-29 | 2012-11-10 | 11.4 days |
|  | 116493_2 | 233 | 2012-11-10 | 2012-11-22 | 12.4 days |
| 116490 (2) | 116490_2 | 229 | 2012-10-31 | 2012-11-15 | 15.0 days |
|  | 116490_3 | 148 | 2012-11-16 | 2012-11-22 | 6.9 days |
| 116488 (1) | 116488_1 | 533 | 2012-10-29 | 2012-11-22 | 24.2 days |
| 116482 (2) | 116482_1 | 121 | 2012-10-29 | 2012-11-07 | 9.0 days |
|  | 116482_2 | 293 | 2012-11-08 | 2012-11-22 | 14.4 days |
| 106384 (3) | 106384_2 | 88 | 2011-11-03 | 2011-11-09 | 5.4 days |
|  | 106384_3 | 204 | 2011-11-09 | 2011-11-23 | 14.3 days |
| 107829 (1) | 107829_1 | 305 | 2011-10-29 | 2011-11-23 | 25.2 days |
| 106389 (1) | 106389_1 | 74 | 2011-10-29 | 2011-11-03 | 5.0 days |
| 106388 (1) | 106388_1 | 481 | 2011-10-29 | 2011-11-23 | 25.2 days |
| 106387 (3) | 106387_1 | 84 | 2011-10-29 | 2011-11-04 | 6.1 days |
|  | 106387_2 | 123 | 2011-11-05 | 2011-11-15 | 10.5 days |
|  | 106387_3 | 113 | 2011-11-16 | 2011-11-23 | 7.6 days |
| 106385 (2) | 106385_1 | 121 | 2011-10-28 | 2011-11-06 | 9.1 days |
|  | 106385_2 | 262 | 2011-11-07 | 2011-11-23 | 16.5 days |
| 116487 (1) | 116487_1 | 448 | 2012-10-28 | 2012-11-22 | 25.8 days |
| 116492 (1) | 116492_10 | 146 | 2012-11-13 | 2012-11-22 | 9.5 days |
| 116491 (1) | 116491_1 | 468 | 2012-10-25 | 2012-11-22 | 28.0 days |
| 116483 (1) | 116483_1 | 399 | 2012-10-25 | 2012-11-22 | 28.2 days |
| 116489 (2) | 116489_1 | 173 | 2012-10-25 | 2012-11-09 | 15.0 days |
|  | 116489_5 | 64 | 2012-11-17 | 2012-11-22 | 5.4 days |
| 116494 (2) | 116494_1 | 91 | 2012-10-23 | 2012-11-02 | 9.4 days |
|  | 116494_2 | 229 | 2012-11-02 | 2012-11-22 | 19.4 days |
| 116495 (1) | 116495_1 | 498 | 2012-10-23 | 2012-11-22 | 30.4 days |
| 116486 (3) | 116486_1 | 75 | 2012-10-24 | 2012-10-29 | 5.7 days |
|  | 116486_4 | 198 | 2012-11-04 | 2012-11-19 | 14.3 days |
|  | 116486_5 | 69 | 2012-11-19 | 2012-11-22 | 3.5 days |
| 116496 (1) | 116496_1 | 588 | 2012-10-23 | 2012-11-22 | 30.2 days |
| 43847 (1) | 43847_1 | 605 | 2010-10-20 | 2010-11-23 | 34.9 days |
| 43864 (1) | 43864_1 | 716 | 2010-10-19 | 2010-11-23 | 35.1 days |
| 43842 (2) | 43842_2 | 210 | 2010-11-09 | 2010-11-23 | 14.4 days |
| 43865 (2) | 43865_1 | 65 | 2010-10-18 | 2010-10-23 | 4.2 days |
|  | 43865_2 | 476 | 2010-10-23 | 2010-11-23 | 31.4 days |
| 43835 (1) | 43835_1 | 743 | 2010-10-17 | 2010-11-23 | 37.9 days |
| 43862 (2) | 43862_1 | 108 | 2010-10-16 | 2010-10-22 | 6.1 days |
|  | 43862_2 | 85 | 2010-10-23 | 2010-10-28 | 4.5 days |
| 43843 (2) | 43843_1 | 162 | 2010-10-16 | 2010-10-25 | 8.3 days |
|  | 43843_2 | 83 | 2010-10-25 | 2010-10-30 | 5.1 days |

**Table A2**. Results from the top three move persistence models of ringed seals in Hudson Bay, ranked by change in AIC. The estimate (Est) and standard error (SE) are presented for scaled parameters and therefore represent relative values instead of raw values.

|  | **Model formula** | **AIC (ΔAIC)** | **Parameter** | **Est** | **SE** | **z-value** | **p-value** |
| --- | --- | --- | --- | --- | --- | --- | --- |
| div2: | ~ diversity + bathy + (1 \| id) | -115846.2 (0) | Intercept | 0.744 | 0.094 | 7.882 | <0.001 |
|  |  |  | diversity | 0.166 | 0.027 | 6.237 | <0.001 |
|  |  |  | bathy | -0.481 | 0.026 | -18.31 | <0.001 |
| fish8: | ~ biomassnsandlance + bathy + (1 \| id) | -115844.1( (2.1) | Intercept | 0.743 | 0.094 | 7.919 | <0.001 |
|  |  |  | biomassnsandlance | 0.163 | 0.027 | 6.001 | <0.001 |
|  |  |  | bathy | -0.467 | 0.026 | -18.18 | <0.001 |
| env2: | ~ bathy + (1 \| id) | -115823.6 (22.6) | Intercept | 0.742 | 0.093 | 7.982 | <0.001 |
|  |  |  | bathy | -0.449 | 0.026 | -17.58 | <0.001 |

diversity = Simpson’s Diversity Index of 8 most important prey species; bathy = bathymetry; biomassnsandlance = biomass of northern sand lance.

**Table A3.** Results from leave-one-out cross validation of move persistence mixed models, for the top selected model (div2), and the third-best model (env2). We removed each individual was iteratively, ran the model at each iteration, and examined the coefficient estimates (estimate and standard error) relative to the full model (i.e., all seals included) coefficient estimates. We report the percent of leave-one-out models where the coefficient estimates fall within the 95% CI of the parameters of the full model. Leave-one-out cross validation for the best non-prey model was ran for comparison to determine if this model performed better than the best-selected model which incorporated modelled prey covariates. The mean estimate (Mean Est) and 95% quantile range (5% and 95%) are presented for scaled parameters and therefore represent relative values instead of raw values. The estimated trend (Est Trend) represents the percentage of cross validation models where the estimated coefficients fall within the 95% CI of the estimates from the final (full) model estimates.

| **Parameter** | **Mean Est** | **5%** | **95%** | **Est Trend** |
| --- | --- | --- | --- | --- |
| **div2: ~diversity + bathy + (1 \| id)** | | | | |
| Intercept | 0.744 | 0.656 | 0.878 | **100%** |
| diversity | 0.166 | 0.152 | 0.206 | **92%** |
| bathy | -0.481 | -0.492 | -0.437 | **96%** |
| **env2: ~bathy + (1 \| id)** | | | | |
| Intercept | 0.742 | 0.660 | 0.874 | **100%** |
| bathy | -0.449 | -0.459 | -0.407 | **94%** |

diversity = Simpson’s Diversity Index of 8 most important prey species; bathy = bathymetry.

**Table A4**. Results from linear mixed effects models between move persistence, northern sand lance biomass, prey diversity, and bathymetry against various dive metrics.

| **Dive metric** | **Est** | **SE** | **p-value** |
| --- | --- | --- | --- |
| **Move persistence (null model) ~** | | | |
| Number of dives | -0.0001 | <0.0001 | 0.717 |
| Maximum depth (m) | 0.0007 | <0.0001 | <0.001 |
| Sum of dives (seconds) | 0.0001 | <0.0001 | <0.001 |
| Mean maximum depth (m) | 0.0007 | <0.0001 | <0.001 |
| Sum depth of all dives (m) | 0.0001 | <0.0001 | <0.001 |
| Sum bottom time (seconds) | 0.0001 | <0.0001 | 0.002 |
| Mean bottom time (seconds) | 0.0001 | <0.0001 | <0.001 |
| **Move persistence (best model: div2) ~** | | | |
| Number of dives | -0.0001 | <0.0001 | 0.257 |
| Maximum depth (m) | 0.0006 | <0.0001 | <0.001 |
| Sum of dives (seconds) | 0.0001 | <0.0001 | <0.001 |
| Mean maximum depth (m) | 0.0007 | <0.0001 | <0.001 |
| Sum depth of all dives (m) | 0.0001 | <0.0001 | <0.001 |
| Sum bottom time (seconds) | 0.0001 | <0.0001 | <0.001 |
| Mean bottom time (seconds) | 0.0001 | <0.0001 | <0.001 |
| **Northern sand lance biomass ~** | | | |
| Number of dives | 0.0001 | <0.0001 | 0.802 |
| Maximum depth (m) | -0.0001 | <0.0001 | 0.223 |
| **Prey diversity ~** | | | |
| Number of dives | <0.0001 | <0.0001 | 0.802 |
| Maximum depth (m) | <0.0001 | <0.0001 | 0.298 |
| **Bathymetry ~** | | | |
| Number of dives | -0.0007 | 0.0004 | 0.713 |
| Maximum depth (m) | -9.2168 | 0.4887 | <0.001 |
| Proportion of depth dove to | 0.094 | 0.0022 | <0.001 |

Est = estimate, SE = standard error.

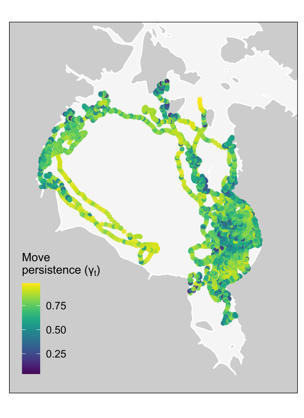

**Figure A1.** Map of seal locations coloured by estimated move persistence (γ_t_) from our best model, which included prey diversity and bathymetry, where low move persistence is indicative of area-restricted search (foraging), and high move persistence is indicative of direct movement (travelling).

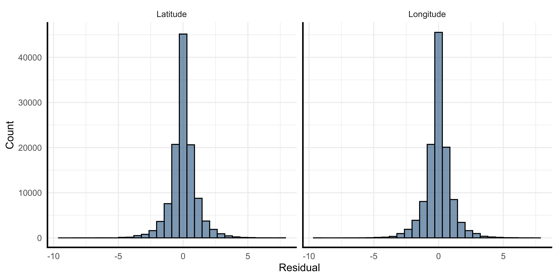

**Figure A2**. Histogram of the one-step-ahead residuals from the best move persistence mixed model, div2, for both the latitude (left) and the longitude (right). The leptokurtic distribution (tails wider than normal distribution) suggests a minor deviation from normality.

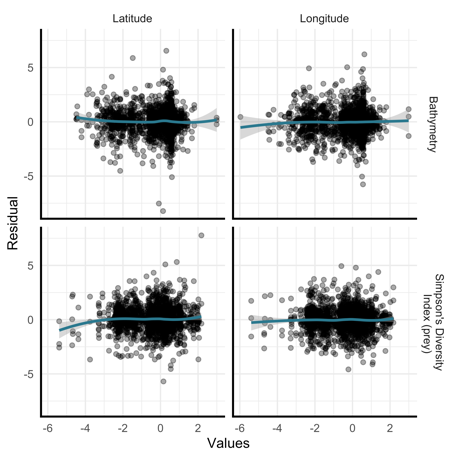

**Figure A3**. Plots of the one-step-ahead residuals from the best move persistence mixed model, div2, for both the latitude (left) and the longitude (right) with each covariate (scaled). There are no obvious patterns between the residuals and covariates, which suggests that the linear assumption is adequate.

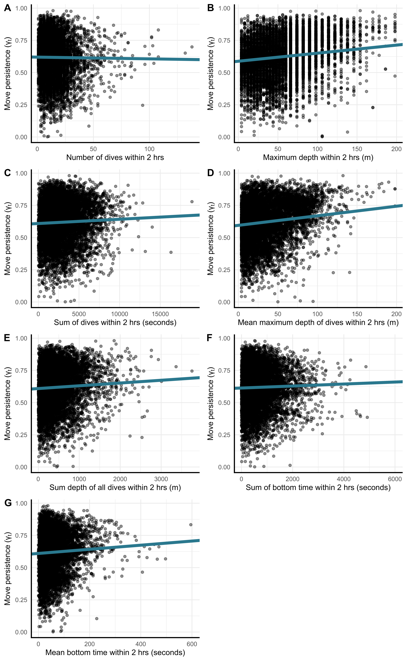

**Figure A4.** Relationships between move persistence (γ_t_ from the best model: div2) and dive metrics from ringed seals in Hudson Bay. Low move persistence is indicative of area restricted search (foraging), and high move persistence is indicative of direct movement (travelling). We expected a negative relationship between move persistence and the dive metrics, where low move persistence (foraging) should correlate with more diving effort.

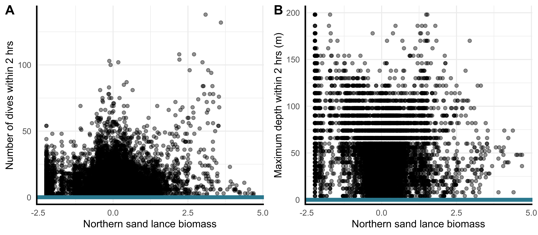

**Figure A5**. Relationships between northern sand lance biomass and dive metrics from ringed seals in Hudson Bay. We expected a positive relationship between prey biomass and the dive metrics.

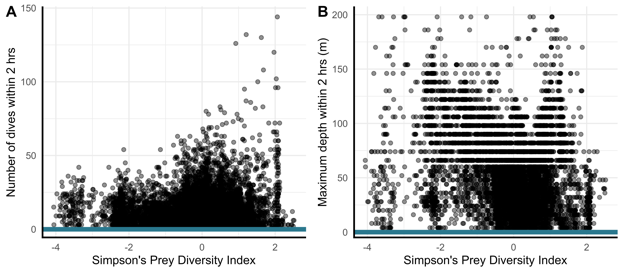

**Figure A6**. Relationships between Simpsons Prey Diversity Index and dive metrics from ringed seals in Hudson Bay. We expected a positive relationship between prey diversity and the dive metrics.

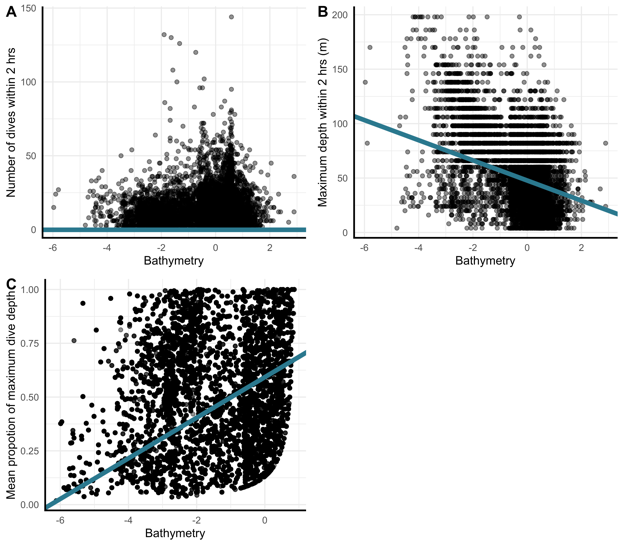

**Figure A7**. Relationships between bathymetry (scaled) and dive metrics from ringed seals in Hudson Bay. We expected seals to dive more frequently in shallower waters, dive deeper in deeper waters, and use a greater proportion of the water column in shallow waters.

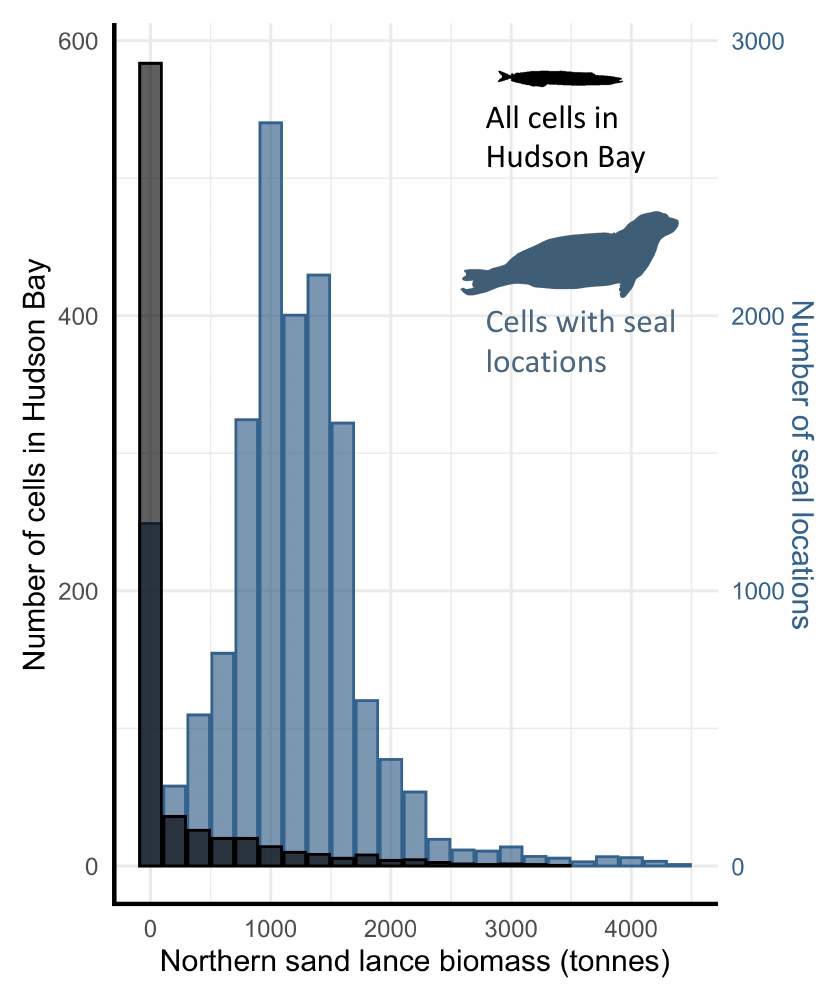

**Figure A8**. Histogram of northern sand lance biomass in the 0.5 ̊ longitude by 0.5 ̊ latitude grid cells of the study area at the mean study year, 2009 (black, left y-axis), and northern sand lance biomass relative to the number of ringed seal locations (blue, right y-axis).
